# Ambient temperature storage in DESS supports molecular studies of benzimidazole resistance from canine hookworm eggs

**DOI:** 10.1101/2025.08.16.670690

**Authors:** Yi-Jou Chen, Vanessa Li, Michelle Suwandy, Ioana Bianca Mitrea, Douglas Hayward, Susan Jaensch, Emily Kate Francis, Jan Šlapeta

## Abstract

Emerging benzimidazole resistance in canine hookworms poses a growing concern for veterinary and public health. Molecular diagnostics targeting β-tubulin gene mutations are essential for resistance surveillance but traditionally rely on refrigerated faecal samples. This study evaluates dimethyl sulfoxide, EDTA, and saturated NaCl (DESS) as a room-temperature preservation medium for canine faeces. Using ITS-2 rDNA and isotype-1 β-tubulin gene (*tubb-1*) amplicon metabarcoding, we compared DNA integrity and diagnostic performance of DESS-preserved samples (4, 28, and 106 days) to refrigerated controls. No significant differences in PCR amplification or sequencing outcomes were observed. DESS enabled reliable detection of hookworm species and resistance-associated SNPs, including F167Y in *tubb-1*, with mutation frequencies consistent across treatments. Therefore, DESS can preserve samples from remote settings without cold chain logistics. Our findings validate DESS as a robust alternative for sample collection for molecular parasitology, facilitating expanded surveillance of anthelmintic resistance in field conditions.

**Highlights:**

1. DESS preserves canine faeces for DNA analysis without refrigeration.
2. Hookworm species and resistance SNPs detected reliably from DESS samples.
3. PCR and NGS results from DESS samples match refrigerated controls.
4. F167Y mutation found in *A. caninum* and *U. stenocephala* samples.
5. DESS enables remote hookworm surveillance without cold chain logistics.

## 1. Introduction

Canine hookworms, *Ancylostoma caninum* and to a lesser extent *Uncinaria stenocephala*, are frequent and impactful gastrointestinal parasites of domestic dogs worldwide (Hawdon and Wise, 2021; Evason et al., 2023). In addition, they are both zoonotic (Bloomquist and Elston, 2024). Hookworm infections pose a serious health concern in both human and veterinary medicine, as they can cause potentially fatal disease in young dogs, and lead to conditions such as cutaneous larval migrans and eosinophilic enteritis in humans (Hawdon & Wise 2021). Treatment of canine hookworm relies on three key anthelmintic classes: benzimidazoles, avermectins and tetrahydropyrimidines (Geary et al., 2025; Sonmez et al., 2025). Benzimidazoles are the most widely accessible drugs for hookworm management in veterinary medicine, and any emerging resistance must be considered within the broader One Health context (Hawdon and Wise, 2021; Evason et al., 2023). Compelling evidence now demonstrates benzimidazole resistance in *A. caninum* infecting dogs in North America and eastern Australia (Venkatesan et al., 2023; Abdullah et al., 2025). Moreover, reports of multi-anthelmintic drug resistant *A. caninum* are increasing in the USA (Jimenez Castro et al., 2019; Kitchen et al., 2019; Geary et al., 2025).

For benzimidazoles, known mutations in the isotype-1 β-tubulin gene (*tubb-1*) lead to drug failure, and screening for their presence and frequency can identify populations and individuals where benzimidazoles are likely to be ineffective or their use is not warranted. For other drug classes, however, no specific molecular tests are currently available (Geary et al., 2025). Screening for hookworm *tubb-1* mutations is currently possible through in house amplicon metabarcoding using Illumina sequencing or even via commercially available diagnostic tests in the USA (Geary et al., 2025). Traditionally, this testing relies on fresh faecal samples, with concentrated eggs subjected to DNA isolation (Venkatesan et al., 2023; Abdullah et al., 2025). To keep hookworm eggs intact for concentration via flotation techniques, faecal samples must be refrigerated to prevent embryonation. In ruminants, faeces with strongyle eggs can be also preserved by vacuum packing with storage at room temperature (Rinaldi et al., 2011). In dog hookworms the refrigeration requirement poses challenges in regions with limited access to refrigeration and creates logistical constraints when cooled samples need to be shipped. To address the need for refrigeration, we explored the use of faecal preservation with dimethyl sulphoxide, EDTA, and saturated NaCl (DESS), which has previously been shown to maintain DNA integrity in nematodes (Yoder et al., 2006; Beknazarova et al., 2017; Gonzálvez et al., 2022). Originally developed to preserve avian tissues at room temperature, DESS is now widely used across diverse specimens for genetic and genomic analyses, including preservation of high-molecular weight DNA, and has been shown to outperform storage in ethanol for DNA preservation (Seutin et al., 1991; Dawson et al., 1998; Yoder et al., 2006; Gaither et al., 2011; Beknazarova et al., 2017; Oosting et al., 2020; Sharpe et al., 2020; Gonzálvez et al., 2022).

This study aimed to evaluate the use of DESS solution for preserving canine faecal samples at ambient temperature for molecular analysis of presumed strongyle/hookworm-like eggs. We applied strongyle species identification using ‘nemabiome’ ITS-2 rDNA amplicon metabarcoding and targeted hookworm beta-tubulin isotype 1 (*tubb-1*) assays to assess whether samples stored in DESS for 4, 28 and 106 days yielded equivalent results to refrigerated canine faecal samples.

## 2. Material and Methods

### 2.1. Samples and samples processing

The use of residual clinical samples was in accordance with the University of Sydney Animal Ethics Committee Protocol (2024/2508). Faecal samples (n=31, Table 1) were provided by registered veterinary practitioners in accordance with the Veterinary Practice and Animal Welfare Acts for the purpose of diagnostics used in decision-making by the veterinary practitioner. Samples were submitted for parasitological diagnostics; refrigerated residual faecal samples that previously returned a positive result for hookworms at the clinical diagnostic laboratory were received for molecular species characterisation. Material used for hookworm species identification and benzimidazole susceptibility, and subsequent results were de-identified with data of samples collection, age, locality (postcode) and breed retained.

**Table 1:** Summary of dog samples with hookworm-like eggs used in this study.

| Identifier | Breed | Age | Date | EPG | Locality | *A. caninum tubb-1* (%) |  | *U. stenocephala tubb-1* (%) |  |
| --- | --- | --- | --- | --- | --- | --- | --- | --- | --- |
|  |  |  |  |  |  | Q134H | F167Y | Q134H | F167Y |
| ACH01* | Staghound cross | 3 yrs | February 2025 | <40 | QLD 4869 | 0, 0, 0 | 0, 0, 0 | n/d | n/d |
| ACH02* | Chihuahua | n/a | November 2024 | 200 | TAS 7000 | n/d | n/d | n/d | n/d |
| ACH03* | Cavoodle | n/a | November 2024 | 3,500 | NSW 2030 | 0, 0, 0 | 0, 0, 0 | n/d | n/d |
| ACH04* | Italian Greyhound | 3 yrs | November 2024 | 400 | NSW 2290 | n/d | n/d | n/d | n/d |
| ACH05* | Greyhound | 5.9 yrs | November 2024 | 240 | NSW 2620 | 0, 48.6, 0 | 0, 0, 0 | n/d | n/d |
| ACH06* | Beaglier | 1.2 yrs | November 2024 | 80 | NSW 2102 | n/d | n/d | n/d | n/d |
| ACH07* | Miniature Fox Terrier | 0.5 yrs | April 2020 | 80 | NSW 2570 | n/d | n/d | n/d | n/d |
| ACH08* | Kelpie | 0.2 yrs | December 2024 | 2,200 | NSW 2312 | 0, 0, 0 | 0, 0, 0 | n/d | n/d |
| ACH09* | Kelpie | 0.2 yrs | December 2024 | 1,600 | NSW 2312 | 0, 0, 0 | 0, 0, 0 | 0, 0, 0 | 44.4, 48.0, 75.3 |
| ACH10* | Greyhound | 2 yrs | February 2025 | 7,160 | ACT 2904 | 0, 3.5, 4.6 | 9.5, 17.4, 14.7 | n/d | n/d |
| ACH11* | Pug | 1 yr | January 2024 | 2,360 | NSW 2594 | 0, 0, 0 | 0, 0, 0 | n/d | n/d |
| ACH12* | Border Collie | 0.1 yr | January 2025 | 7,680 | QLD 4067 | 0, 0, 0 | 0, 0, 0 | n/d | n/d |
| ACH13* | Miniature Schnauzer | 11 yrs | January 2025 | 1,560 | NSW 2444 | 0, 0, 0 | 0, 0, 0 | n/d | n/d |
| ACH14* | Jack Russell Terrier | 5 yrs | December 2024 | 4,240 | NSW 2321 | 0, 0, 0 | 0, 0, 0 | n/d | n/d |
| ACH15* | Fox Hound | 4 yrs | December 2024 | 23,280 | QLD 4067 | 0, 0, 0 | 0, 0, 0 | n/d | n/d |
| ACH16 | Greyhound | 4 yrs | February 2025 | 160 | NSW 2795 | 0 | 18.5 | n/d | n/d |
| ACH17 | Labrador Retriever | 2 yrs | February 2025 | 125 | NSW 2711 | 0 | 0 | 0 | 0 |
| ACH18 | Doberman | 0.6 yr | March 2025 | 750 | QLD 4390 | 0 | 0 | n/d | n/d |
| ACH19 | Shar-Pei | 6.6 yrs | April 2025 | 400 | NSW 2460 | 2.7 | 95.5 | n/d | n/d |
| ACH20 | Greyhound | 2 yrs | April 2025 | 560 | NSW 2430 | n/d | n/d | n/d | n/d |
| ACH21 | Griffan Bruxelloxs | 2 yrs | March 2025 | 12,040 | QLD 4124 | 0 | 0 | n/d | n/d |
| ACH22 | Staffordshire Bull Terrier | 0.6 yrs | August 2024 | 440 | NSW 2621 | n/d | n/d | n/d | n/d |
| ACH23 | French Bulldog | 1 yr | September 2024 | 1,680 | NSW 2711 | 0 | 16.4 | n/d | n/d |
| ACH24 | Greyhound | 2 yrs | July 2024 | 20,000 | NSW 2460 | 0 | 0 | n/d | n/d |
| ACH25 | Greyhound | 0.6 yr | May 2024 | 240 | QLD 4570 | 2.1 | 53.4 | n/d | n/d |
| ACH26 | Greyhound | 1.6 yr | July 2024 | <40 | NSW 2590 | 0 | 0 | n/d | n/d |
| ACH27 | Labrador Retriever | 3.5 yrs | July 2024 | <40 | NSW 2794 | n/d | n/d | n/d | n/d |
| ACH28 | Greyhound | 1.9 yrs | May 2024 | 40 | NSW 2594 | 0 | 0 | n/d | n/d |
| ACH29 | Greyhound | 3.9 yrs | May 2024 | <40 | NSW 2460 | 20.5 | 10.6 | n/d | n/d |
| ACH30 | Bull Arab cross | 0.3 yr | October 2024 | 600 | NSW 2037 | n/d | n/d | 0 | 3.7 |
| ACH31 | Greyhound | adult | September 2024 | n/a | NSW 2460 | 0 | 0 | n/d | n/d |

\*, samples processed as raw concentrated, DESS-preserved for 4 days and DESS-preserved for 28 days; n/a, not known; EPG, strongyle hookworm-like eggs per gram; QLD, Queensland, Australia; NSW, New South Wales, Australia; TAS, Tasmania, Australia; ACT, Australian Capital Territory, Australia; n/d, not detected.

Refrigerated samples received from diagnostic laboratories were partitioned into (i) 3 g for faecal egg counting and egg concentration, and (ii) 3 g for preservation in DESS. Faecal egg counts and concentrations were performed with saturated salt (NaCl, specific gravity = 1.21) solution using Whitlock Universal McMaster 4 chamber worm egg counting slide at the University of Sydney and eggs concentrated and collected as previously described from the 3 g of faeces via centrifugal flotation (800×g) in 50 mL Falcon-type plastic tube in saturated NaCl (Abdullah et al., 2025). Faecal samples (3 g) were homogenised in 50 mL of NaCl solution and centrifuged at 800 × g for 5 minutes. The upper 10 mL of the supernatant was carefully transferred to a clean 50 mL Falcon-type plastic tube, topped up with distilled water, and centrifuged again under the same conditions. Following centrifugation, the supernatant was aspirated and discarded. The resulting pellet was transferred to a 1.5 mL microcentrifuge tube and washed with distilled water. The limit of detection was 40 eggs per gram and the resultant concentrated strongyle/hookworm-like egg pellets were stored in 1.5 mL tubes at -20 °C until further use or immediately processed for DNA isolation. DESS was prepared according Seutin et al. (1991). We used the following chemicals: dimethyl sulfoxide (DMSO, cat. no. 2225, Ajax Finechem), EDTA disodium salt (cat.no. 180, Ajax Finechem), sodium chloride (NaCl analytical grade, cat. no. 465, Ajax Finechem) and sodium hydroxide (NaOH, analytical grade, cat. no. 482, Ajax Finechem). DESS is 0.25M disodium EDTA pH 8.0 (pH adjusted using NaOH), 20% DMSO and NaCl added to saturation. DESS was aliquoted into 15 mL Falcon-type plastic tubes each containing 10 mL of DESS. Faeces to be preserved in DESS were transferred (3 g) into the 15 mL Falcon-type plastic tube with 10 mL of DESS, vigorously shaken and stored at room temperature (23-24 °C), aliquot (∼1.5 mL of DESS with faeces) was subsampled on day 4, day 28 and day 106.

### 2.2. DNA isolation

Genomic DNA was isolated using Isolate II Fecal DNA Kit (Meridian Bioscience, Australia). DESS-preserved samples were homogenised by inverting the 15 mL tube with faeces in DESS multiple times, then immediately a ∼1.5 mL aliquot was aspirated and transferred into 2 ml bead-beating tube from Isolate II Fecal DNA Kit (Meridian Bioscience, Australia) and centrifuged at 3,000×g for 5 minutes. The supernatant was discarded and pellet flooded with 750 µL of lysis buffer from the DNA kit. The pellet with lysis buffer was homogenised using a high-speed benchtop homogeniser, FastPrep-24 (MP Biomedicals, Australia), for 40 s at 6.0 m/s. For concentrated strongyle/hookworm-like eggs in 1.5 mL tube, they were first flooded with 750 µL of lysis buffer and homogenised by repeated aspiration with a pipette, then all the contents were transferred into bead-beating tube from Isolate II Fecal DNA Kit (Meridian Bioscience, Australia) and the eggs disrupted and homogenised using FastPrep-24 using setting as described above. Blank samples that only contained the lysis buffer we included throughout the process. Purified genomic DNA from all samples and blanks were eluted into 100 µL of elution buffer (10 mM Tris-Cl, pH = 8.5) and stored at −20 °C until further use.

### 2.3. Amplification and comparison

All DNA samples were subjected to two independent real-time PCR amplifications. ITS-2 rDNA (ITS-2) was amplified with the NC-1 and NC-2 primers designed by Gasser et al. (1993), and isotype-1 β-tubulin region BZ167 was amplified using primers developed by Jimenez Castro et al. Jimenez Castro et al. (2019), which include key diagnostic SNPs conferring benzimidazole resistance. All primers were adapted for Illumina metabarcoding using Next Generation Sequencing (NGS) as previously described (Stocker et al., 2023; Abdullah et al., 2025). All PCR reactions utilised SYBR-chemistry using SensiFAST™SYBR® No-ROX mix (Meridian Bioscience, Australia). All PCR reactions were prepared using a Myra robotic liquid handling system (Bio Molecular Systems, Australia). Each real-time PCR mix (25 μL) included 2 μL of template DNA (neat; from both the concentrated material and the DESS preserved material) and primers at final concentrations of 400 nM. Reactions were run on a CFX Opus 96 Real-Time PCR System with corresponding CFX Manager software (BioRad, Australia) to obtain cycle threshold values (Ct). The protocol involved an initial denaturation at 95 °C for 3 min, followed by up to 35 cycles of 95 °C for 5 s, 57 °C for 15 s, 72 °C for 15 s, and a final melt curve analysis. Each run included a no-template control (ddH_2_O) to monitor for potential contamination. Ct>35 samples were recoded as 35. Ct values for all samples were tabulated and analysed in GrapPad Prism v10.4.1 (GraphPad Software). Data were analysed separately for ITS-2 and BZ167 assays. Data passed tests for Gaussian normal distribution (D’Agostino-Pearson and Shapiro-Wilk tests). Paired t-test was run for raw concentrated egg Ct compared to Ct from samples stored in DESS for 4 days. Repeated measure (RM) one-way ANOVA with Geisser-Greenhouse correction and Tukey’s multiple comparison test was run for all treatments (raw concentrated, DESS 4 days, DESS 28 days, DESS 106 days) per assay. RM one-way ANOVA was rerun within the data that has potentially ambiguous data coded as Ct=35. The alfa (α) was set at 0.05.

### 2.4. Next generation sequencing and data processing

Amplicons (ITS-2 and BZ167) for raw concentrated, DESS 4 days and DESS 28 days as described above were prepared in 96 well plate format and submitted for indexing, purification and amplicon sequencing. In addition, for raw concentrated samples BZ200 amplicon was amplified as described above using primers developed by Jimenez Castro et al. (2019) as implemented previously (Stocker et al., 2023; Abdullah et al., 2025). The next generation sequencing was performed at the Ramaciotti Centre for Genomics, University of New South Wales, Sydney, Australia and sequenced on MiSeq i100 Plus 25M (300PE, Ilumina) and MiSeq (250PE, Illumina). De-multiplexed FastQ files were generated via BaseSpace (Illumina).

All FastQ files were processed through R package ‘dada2’ v1.26.0 (Callahan et al., 2016) in R v4.2.2 (R Core Team, 2022), to obtain amplicon sequence variants (ASV) and their abundance per sequenced amplicon using pipeline developed previously (Abdullah et al., 2025). The pipeline includes, primer removal with ‘cutadapt’ v 4.0 (Martin, 2011), followed by ‘dada2’ filtering, error estimation and denoising, before paired-end reads were merged to reconstruct the full ASV and removal of chimeras. The assignment of species based on ITS-2 ASVs was carried out using two complimentary approaches. We used IdTaxa from ‘DECIPHER’ package v3.2.0 (Murali et al., 2018) using ITS-2 databased from nemabiome.ca (Workentine et al., 2020) and assignTaxonomy from ‘dada2’ using in house hookworm database. All ASVs were further imported into CLC Main Workbench v25.0.1 (Qiagen Australia) and top blastn hits against ‘nr’ database at NCBI was identified. ASVs multiple sequence alignment was subject to phylogenetic reconstruction with reference ITS-2 hookworm sequences, where species identity was confirmed by manual assignment of ASV to a species based on position on the phylogenetic tree relative to the reference ITS-2 sequences. For the *tubb-1* data (BZ167, BZ200), the assignment to species was carried as previously described (Abdullah et al., 2025). Briefly, all ASVs assignment was verified in CLC Main Workbench v25.0.1 against curated set of *tubb-1* reference sequences from *A. caninum*, *A. ceylanicum*, *A. duodenale* and *Necator americanus* from ‘WormBase ParaSite’ (Howe et al., 2017; Bolt et al., 2018; Harris et al., 2020) and *U. stenocephala* (Stocker and Šlapeta, 2025). The sequence comparison enabled us to identify the exon/intron boundary and residues that code for susceptibility to benzimidazoles, amino acids amino acids Q134, F167, E198 and F200 (Venkatesan et al., 2023). ASVs were then pooled according to species and raw read counts were converted into relative abundance (percentages). Any species read counts with less <25 reads were disregarded as spurious. Proportions of SNP for amino acids amino acids Q134, F167, E198 and F200 were calculated per species for each of the amplicons of *tubb-1* (BZ167, BZ200).

### 2.5. Data availability

The metadata and intermediate files for all samples are available at LabArchives (https://dx.doi.org/10.25833/pxca-6w16). Raw FastQ sequence data was deposited at SRA NCBI BioProject: PRJNA1300611.

## 3. Results

### 3.1. Canine faecal samples stored in DESS solution at ambient temperature are non-inferior to refrigerated samples using ITS2 rDNA assay targeting strongyles and *tubb-1* of hookworms

Residual diagnostic canine faecal samples (n=31), previously determined to be positive for hookworm-like eggs (strongyle-type eggs) using faecal flotation technique, were used throughout (Table 1). These samples were processed for molecular determination of the identity of the hookworm-like eggs (ITS-2 assay) and potential carriage of the *tubb-1* alleles conferring benzimidazole resistance in hookworms. Samples (n=15) were split into two treatment groups: (i) for processing using centrifugal flotation to concentrate the hookworm-like eggs and immediate DNA isolation, and (ii) for preserving the raw faecal samples in DESS solution and kept at ambient temperature. From the DESS-preserved samples, an aliquot was collected and DNA isolated on day 4, day 28 and day 106.

Amplification of ITS-2 from concentrated eggs’ DNA (n=15) and DESS-preserved samples DNA after 4 days preservation at ambient temperature (n=15) yielded a mean Ct of 23.41 (min. 15.67, max. 34.00, SD 5.95) and a mean Ct of 22.72 (min. 15.07, max. 34.81, SD 4.72), respectively. There was no significant difference between the two sets of Ct in a paired t-test (two-tailed P=0.64, t=0.48, df=14). Testing samples at 28 days and 106 days stored in DESS resulted in no significant Ct difference between the concentrated eggs DNA and between any of the DESS-preserved samples DNA, using RM one-way ANOVA with Geisser-Greenhouse correction and Tukey’s multiple comparison test. Removing ACH02, ACH04, ACH06 and ACH07 (see Section 3.2; Table 1) had no effect on the calculation, and the resulting comparisons between sample (n=11) remained non-significant (Figure 1A).

**Figure 1.**
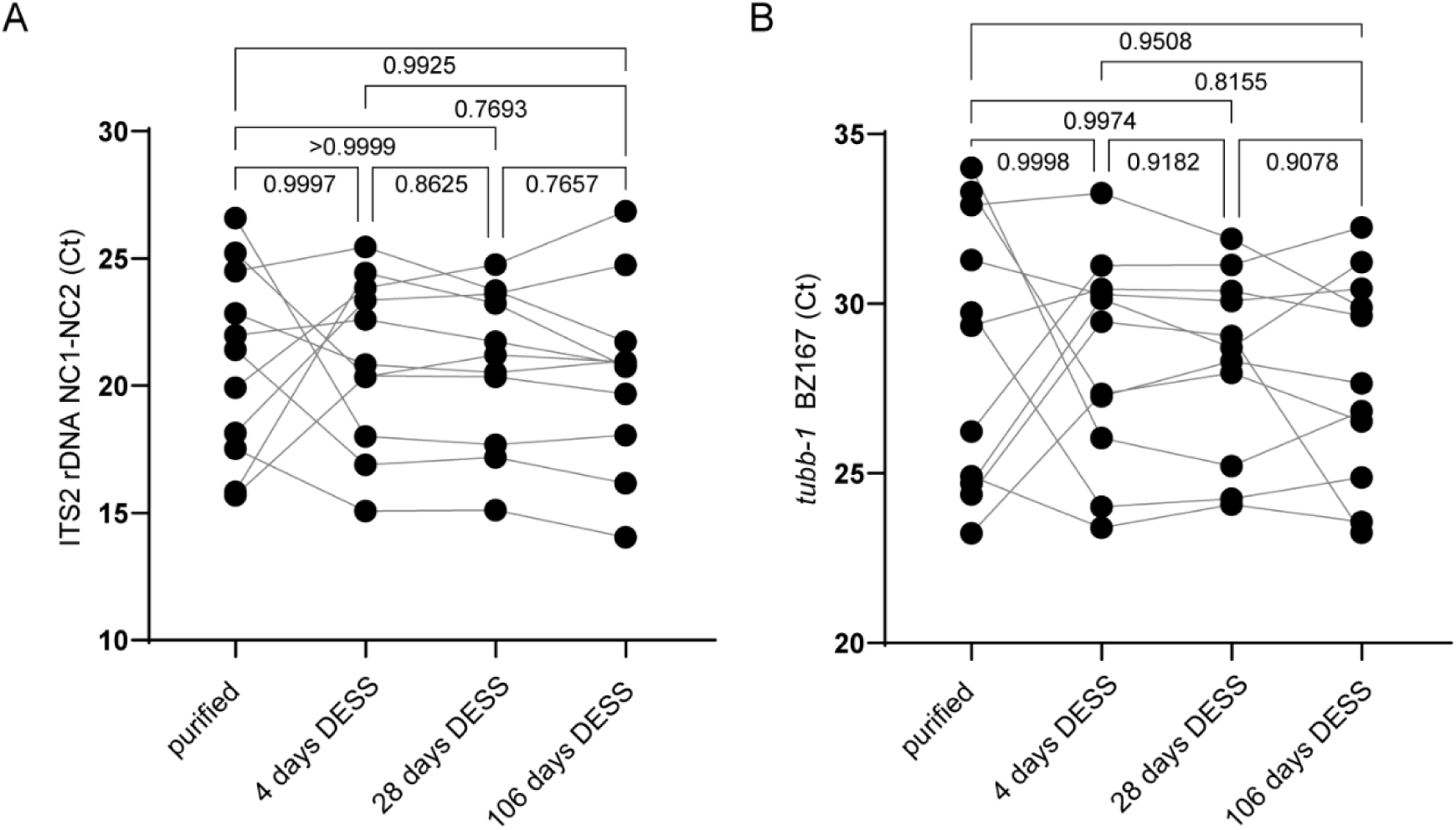
Real-time PCR amplifications of (A) internal transcribed spacer (ITS)-2 rDNA and (B) isotype-1 β-tubulin gene (*tubb-1*) from dog faecal samples. Samples with hookworm-like eggs were amplified and cycle threshold (Ct) was recoded. The *tubb-1* amplicon BZ167 spans the mutations responsible for Q134H and F167Y amino acid changes. Only samples that were confirmed to contain hookworm species are shown. Each point represents an individual Ct value. Each sample was processed as either concentrated eggs (purified eggs) or preserved in DESS for different length of time - 4 days, 28 days and 106 days. Each faecal sample is connected by thin line. Repeated measure (RM) one-way ANOVA with Geisser-Greenhouse correction and Tukey’s multiple comparison test demonstrated no significant difference between the treatments (P>0.05).

Amplification of BZ167-product from *tubb-1* from concentrated eggs’ DNA (n=15) and DESS-preserved samples DNA after 4 days preservation at ambient temperature (n=15) yielded a mean Ct of 28.83 (min. 23.23, max. 34.69, SD 3.84) and a mean Ct of 29.00 (min. 23.40, max. 34.06, SD 3.13), respectively. There was no significant difference between the two sets of Ct in a paired t-test (two-tailed P=0.88, t=0.16, df=14). Similarly, Ct showed no significant differences between concentrated egg DNA and DESS-preserved samples at 28 and 106 days, based on RM one-way ANOVA with Geisser-Greenhouse correction and Tukey’s test. Removing ACH02, ACH04, ACH06, and ACH07 did not affect the outcome; comparisons among the remaining samples (n = 11) remained non-significant (Figure 1B).

### 3.2. Amplicon metabarcoding from refrigerated canine faecal samples is non-inferior in detecting hookworm species and hookworm *tubb-1* SNP conferring benzimidazole resistance

The amplions from the ITS-2 assay were subject to deep-sequening (‘nemabiome’ amplicon metabarcoing). For the 15 samples treated with DESS, we sequenced both the ampliocns from the concentrated hookworm-like eggs DNA as well as amplicons from DNA isolated from DESS stored samples for 4 and 28 days (Figure 2). In addition, we re-isolated eggs from ACH06, isolated DNA and amplified ITS-2. An additional 16 suspect hookworm positive samples were included (Figure 2). After quality control and taxonomic assignment, all but five samples yealded >2,000 good quality sequence reads belonging to strongyles (Figure 2A). No strongyle reads were produced for ACH04 samples and for one of the ACH06 samples. In addition, ACH20 has only 115 strongyle reads. For the remainig samples sequening depth ranged from 83,594 to 5,287 reads (Figure 2A).

**Figure 2.**
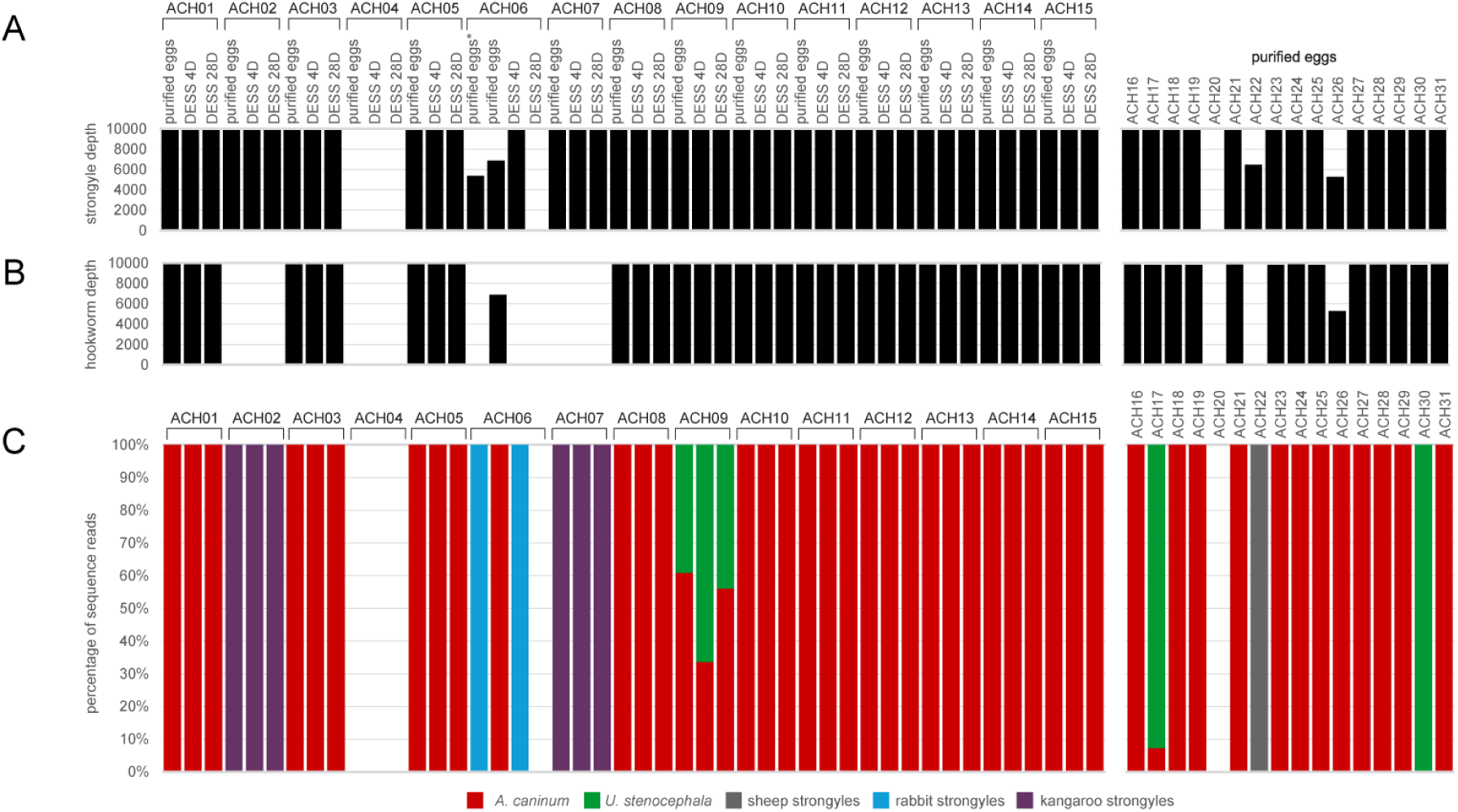
Internal transcribed spacer (ITS)-2 rDNA molecular profiling of dog faecal samples. Sampled with hookworm-like eggs were profiled using metabarcoded deep Illumina amplicon sequencing at ITS-2 rDNA. ACH1-15 have been profiled as concentrated eggs (purified eggs), faeces preserved in DESS for 4 days (DESS 4D) and 28 days (DESS 28D). ACH06 was reisolated as purified eggs and reamplified/re-sequenced (*). Samples ACH16-31 were only processed as concentrated purified eggs. (A) The bar chart representing the number of sequencing reads belonging to strongyle ITS-2 sequences for each sample, the maximum upper scale is set to show 10,000 reads. (B) The bar chart showing sequencing reads belonging to hookworms (Ancylostomatidae) ITS-2 sequences for each sample, the maximum upper scale is set to show 10,000 reads. (C) Percentage of strongyle species within each sample (depicted as proportion). Amplicon sequence variants (ASVs) were pooled according to species of nematodes. Proportion of each hookworm species (*Ancylostoma caninum*, *Uncinaria stenocephala*) and spurious species is colour coded in the stacked bar chart. Samples are vertically aligned across the bar charts.

Considering only hookworm taxonomic asignment based on ITS-2, 11 of the 15 samples had hookworm sequence reads across all the treatments (eggs purification and DESS preseravation) (Figure 2B). Only *A. caninum* or *U. stenocephala* hookworm sequnces were detected (Figure 2C). The ACH09 was mixed infection detected in all the treatmnets (eggs purification and DESS preseravation for 4 days and 28 days). The remainig ten samples had only *A. caninum* sequences consistently detected in all treatments. For the remainig non-hookworm or variable samples, sequences belonged to kangaroo, rabbit or sheep strongyle nametodes. None of the ACH02 and ACH07 samples yealded hookworom reads, instead they were assigned to varing species and genera of strongyles from kangaroos (*Rugopharynx, Monilonema, Cloacina, Cyclostrongylus, Parazoniolaimus, Arundelia, Hypodontus, Wallabinema, Labiomultiplex*). Sample ACH06 yielded variable result, besides one sample not yielding any reads, it either had reads belonging to (*Graphidium strigosum*, *Trichostrongylus retortaeformis*) or hookworm (*A. caninum*); we re-purified eggs, re-isolated DNA and re-amplification yielded reads belonging to rabbit nematodes similarly to those from DESS-preserved aliquot. For ACH20, there were only 46/115 reads belonging to hookworm (*A. caninum*), the remaining reads matched sheep nematode ITS-2 sequences.

We then inquired, if the *tubb-1* BZ167 amplicons were suitable for identification of hookworm species and SNPs coding for benzimidazole resistance (mutations Q134H and F167Y), we processed the amplicons for amplicon metabarcoding and deep-sequencing. Using hookworm *tubb-1* reads at BZ167 region, the results of samples (n=15) were compared between concentrated hookworm-like eggs and those preserved in DESS corroborated the ITS2 rDNA results. Samples ACH02, ACH04, ACH06 and ACH07 did not contain any hookworm *tubb-1* sequence reads, although the samples produced sequencing reads not matching any known targets (Figure 3A). The remaining 11 samples had >2,000 sequencing reads that matched *tubb-1* of hookworms regardless of the treatment (Figure 3B). Similarly for samples ACH16-ACH31, two samples (ACH20 and ACH27) did not yield any hookworm *tubb-1* sequences, and ACH20 was excluded because it yielded only 275 reads (<2,000 threshold). In total, there were 24 samples for which *tubb-1* (BZ167 amplicon) revealed hookworms; 21 with only *A. caninum tubb-1*, one with only *U. stenocephala tubb-1* and two with *tubb-1* from both species (Figure 3C). For the co-infection sample ACH09, there were 26.3% sequencing reads belonging to *U. stenocephala tubb-1* from the concentrated eggs samples, while 53.1% and 31.1% reads belonged to *U. stenocephala tubb-1* in the samples preserved in DESS for 4 days and 28 days, respectively (Figure 3C). Sample ACH17 had 91.6% reads that belonged to *U. stenocephala tubb-1*.

**Figure 3.**
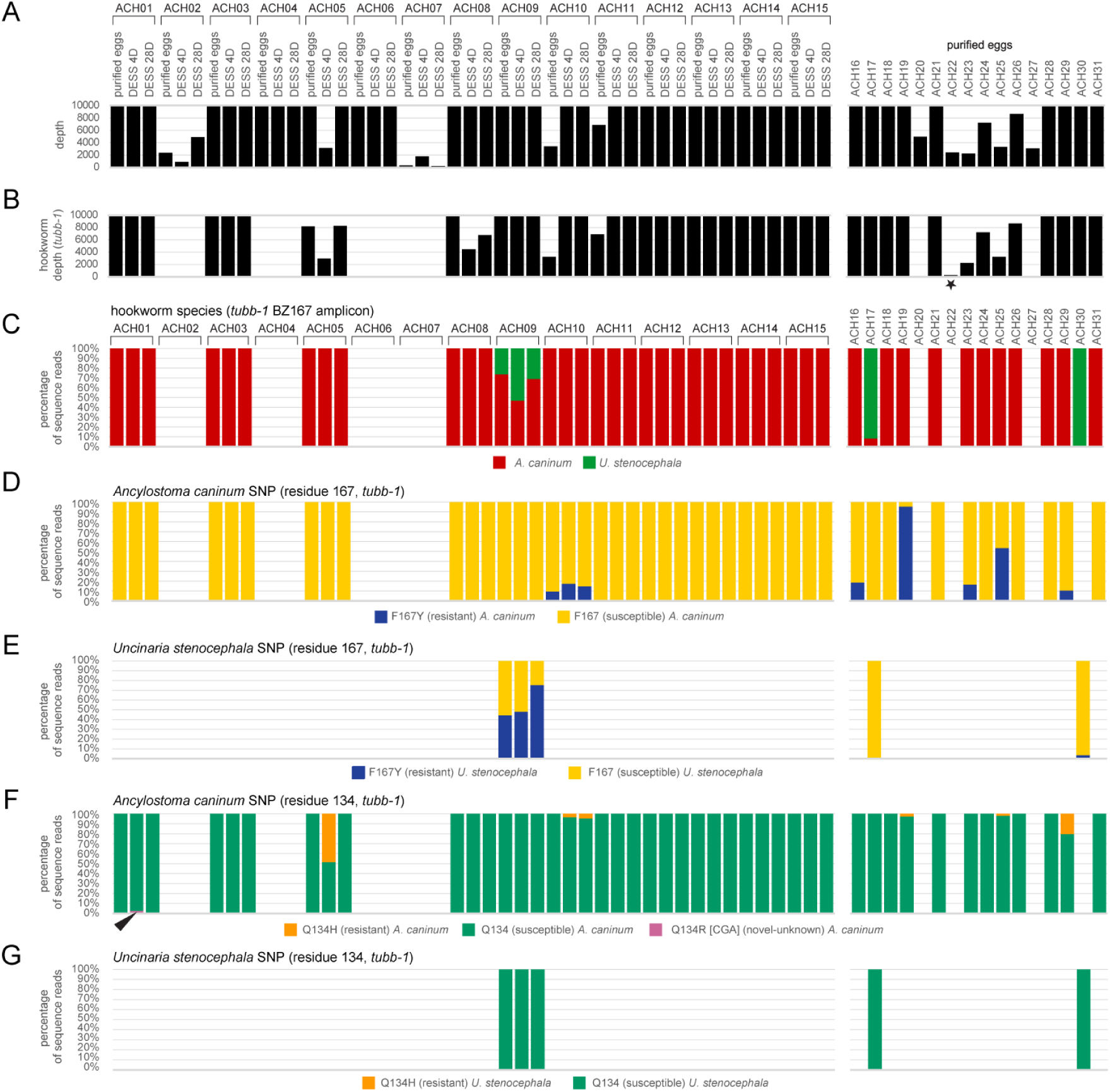
Isotype-1 β-tubulin gene (*tubb-1*) molecular profile of dog faecal samples. Sampled with hookworm-like eggs were profiled using metabarcoded deep Illumina amplicon sequencing at *tubb-1* amplicon BZ167 that spans the mutations responsible for Q134H and F167Y amino acid changes. ACH1-15 have been profiled as concentrated eggs (purified eggs), faeces preserved in DESS for 4 days (DESS 4D) and 28 days (DESS 28D). Samples ACH16-31 were only processed as concentrated purified eggs. (A) The bar chart representing the total number of sequencing reads for each sample, the maximum upper scale is set to show 10,000 reads. (B) The bar chart showing sequencing reads belonging to hookworms (Ancylostomatidae) *tubb-1* sequences for each sample, the maximum upper scale is set to show 10,000 reads. Sample depicted with * (star) had <2,000 reads and was excluded. (C) Percentage of strongyle species within each sample (depicted as proportion) based on the *tubb-1* sequence data. Amplicon sequence variants (ASVs) were pooled according to species of nematodes. Proportion of each hookworm species (*Ancylostoma caninum*, *Uncinaria stenocephala*) is colour coded in the stacked bar chart. (D) Percentage of SNP (single nucleotide polymorphism) that codes for the mutation F167Y in *tubb-1* of *A. caninum*. (E) Percentage of SNP that codes for the mutation F167Y in *tubb-1* of *U. stenocephala*. (F) Percentage of SNP (single nucleotide polymorphism) that codes for the mutation Q134H in *tubb-1* of *A. caninum*. An alternative, novel, mutation leading to Q134R is also shown (arrowhead). (G) No SNP that codes for the mutation Q134H in *tubb-1* of *U. stenocephala*. Samples are vertically aligned across the bar charts.

Mutation at *tubb-1* coding for amino acid change F167Y was present in 26% (6/23) and 33% (2/3) of the samples with *A. caninum* and *U. stenocephala* using the samples from concentrated eggs, respectively (Figure 3D). One of the six *A. caninum* positive samples, ACH10 with 9.5% of F167Y, was part of the DESS treatment; ACH10 was consistently showing F167Y mutation in the DESS-preserved samples for 4 and 28 days (17.4% and 14.7%). One of the two *U. stenocephala* positive samples, ACH10 with 44.4% F167Y mutation, was part of the DESS treatment and consistently showed F167Y mutation allele on day 4 and 28 (48.0% and 75.3%). ACH30 had 3.7% of F167Y mutation allele (Figure 3E). Only one out of the six samples had >70% of the tubb-1 *A. caninum* F167Y (95.5%; ACH19, Table 1) (Figure 3D). A mutation in the *tubb-1* gene resulting in the amino acid substitution Q134H was detected in multiple *A. caninum* DNA samples but was absent in *Uncinaria stenocephala* DNA (Figure 3F). Additionally, a Q134R substitution of unknown functional significance was identified in one of the *A. caninum* samples (ACH01), with a variant allele frequency of 2.6% (Figure 3F).

Dogs that were found positive for *A. caninum* were predominantly greyhounds (39%, 9/23) followed by kelpies (2/23), while the remaining samples came from individual dog breeds (Table 1). Four greyhounds (4/9), one Shar-Pei and one French Bulldog had the *A. caninum tubb-1* mutation F167Y, with the Shar-Pei (ACH19) having the highest percentage of the mutation (95.5%), followed by a greyhound (ACH25, 53.4%). A Kelpie, Labrador Retriever and Bull Arab cross were the breeds that were infected with *U. stenocephala*, the Kelpie (ACH09) sample had the highest percentage (44-75%) of the *U. stenocephala tubb-1* mutation F167Y (Table 1).

All samples screened at *tubb-1* BZ200 amplicons revealed E198 for both *A. caninum* and *U. stenocephala tubb-1*. The F200 residue was conserved across all *U. stenocephala tubb-1*, and majority of *A. caninum tubb-1*, with the exception of rare mutation in F200S (2.8%) in sample ACH18 and F200L (0.1%) in ACH10.

## 4. Discussion

Ensuring optimal data quality in parasitological studies requires careful evaluation of preservation methods. Proper collection and storage are critical for maintaining sample integrity prior to laboratory analysis, yet they often demand considerable effort from field personnel and involve complex logistics to transport samples under suitable conditions. This study demonstrates that preserving canine faecal samples in DESS performs comparably to refrigeration, eliminating the need for a cold chain or reliance on ethanol or proprietary preservatives. Samples stored in DESS remained fully compatible with standard DNA-based molecular workflows, including PCR amplification and NGS-based amplicon sequencing, supporting both hookworm species identification and detection of benzimidazole-resistance associated SNPs.

Using repeated measures one-way ANOVA, we found no significant difference between concentrated egg samples and faecal samples stored in DESS for up to 106 days. DESS preserves DNA by deactivating metal-dependent enzymes (e.g., DNases) via EDTA, with this effect enhanced by the alkaline pH (8) created by NaOH, which dissolves EDTA and provides a buffering capacity. The solution is saturated with sodium chloride (NaCl) to stabilise DNA, and contains DMSO which facilitates the transport of compounds (e.g., EDTA and NaCl) across cell membranes while also acting as a cryoprotectant (Seutin et al., 1991). DESS has been shown to perform well across a broad temperature range, including room temperature. Theoretically, we expected higher Ct values for DESS-preserved samples compared to concentrated samples, as the latter contained the equivalent of 3 g of eggs, whereas only one-tenth of that material was used for the DESS-preserved samples. However, the observed non-inferiority may be explained by material loss during the egg concentration process using saturated salt (Zajac and Conboy, 2011). This represents an additional benefit of DESS, as it suggests egg concentration may not be necessary, potentially streamlining diagnostic workflows using sample volumes similar to those in this study. Although the tested samples had relatively high egg counts (mean: 4,896 eggs per gram; range: <40 to 23,280), samples with lower counts (<500 eggs per gram; ACH01, ACH05) performed comparably to those with higher counts. Nonetheless, caution is warranted when interpreting results from low egg count samples, as insufficient sampling could lead to overinterpretation. Sample-to-sample variability or uneven egg distribution in faecal aliquots was evident in mixed infection samples (ACH09, ACH10). While no studies have investigated the spatial distribution of hookworm eggs in canine faeces, human studies have shown no clear spatial pattern for hookworm and *Schistosoma mansoni* eggs in faecal samples (Krauth et al., 2012). An alternative explanation is that eggs preserved in DESS may continue to embryonate, leading to an increase in cell number and, consequently, a higher number of genomic copies. Similar observations were made with ruminant strongyle eggs preserved in lysis buffer, where embryonation was visually evident and more pronounced compared to eggs stored in ethanol (Francis and Šlapeta, 2022).

The presence of *tubb-1* mutations in hookworm eggs from canine faecal samples does not necessarily equate to clinical drug failure. However, >75% of the F167Y mutation in *tubb-1* has been correlated with in vitro benzimidazole resistance in the Egg Hatch Assay, and can be considered the threshold for predicting clinical resistance based on this mutation alone (McKean et al., 2024; Abdullah et al., 2025). One of our samples, from a Shar-Pei (ACH19), carried 95% of the F167Y mutation, suggesting likely clinical resistance to benzimidazole. Samples with <75% of the F167Y mutation are less likely to lead to observable treatment failure, though continued benzimidazole use may select for this resistant genotype within the hookworm population. In this context, attention should also be given to additional mutations such as Q134H in *tubb-1*, which has been shown to confer benzimidazole resistance even in the absence - or at low frequency - of the F167Y mutation in *A. caninum* (Venkatesan et al., 2023).

Consistent with Australian findings from Abdullah et al. (2025), greyhounds were the most frequently hookworm-positive breed in our study. In North America, analysis of faecal samples from 17,671,724 individual dogs revealed that greyhounds had a significantly higher risk of hookworm infection detected via faecal flotation (odds ratio = 15.3) (Burton et al., 2024). Abdullah et al. (2025) also reported a crossbreed dog infected with *Ancylostoma caninum* carrying >75% of the F167Y mutation, adding to evidence that benzimidazole-resistant *A. caninum* is not restricted to greyhounds in Australia. Although resistance was initially regarded as a breed-specific issue, it now appears to have spread beyond greyhounds (Burton et al., 2024; Geary et al., 2025; Jimenez Castro et al., 2025). A recent North American survey of diagnostic samples found that greyhounds were only the fourth most frequently infected breed with the *tubb-1* F167Y mutation, with poodles, Bernese mountain dogs, and cocker spaniels showing even higher prevalence (Leutenegger et al., 2024; Jimenez Castro et al., 2025).

The *tubb-1* F167Y allele has recently been reported in *U. stenocephala* in Australia, providing an early warning of potential benzimidazole resistance (Abdullah et al., 2025). The allele was detected in a 10-week-old Labrador Retriever with an egg count of 200 (TS024 in Abdullah et al., 2025), where 50.7% of the eggs carried the F167Y mutation. In this study, we report two additional *U. stenocephala* samples carrying the F167Y allele: one from a 10-week-old Kelpie with a mutation frequency between 44–75%, and another with a low frequency (<5%). To date, six *U. stenocephala*-positive samples from southeastern Australia (New South Wales and the Australian Capital Territory) and three from New Zealand have been screened for the F167Y allele (this study; Stocker et al., 2023; Abdullah et al., 2025). In contrast, reanalysis of a nationwide study in the USA by Venkatesan et al. (2023) identified *U. stenocephala* in two individual dog faecal samples and four pooled samples, but all carried the benzimidazole-susceptible F167 allele (Stocker et al., 2024). No studies have yet investigated *U. stenocephala* in dogs from regions where it is common, such as Canada and continental Europe. The presence of the *tubb-1* F167Y allele in *U. stenocephala* is a strong indicator of potential clinical drug failure. Establishing in vivo and in vitro evidence that benzimidazoles fail to clear *U. stenocephala* infections carrying the F167Y allele is now urgently needed.

## Conclusions

This study confirms that DESS is a robust and practical preservation medium for canine faecal samples used in molecular parasitology. It maintains DNA integrity over extended periods without refrigeration, supports PCR and NGS workflows, and enables reliable detection of hookworm species (*A. caninum* and *U. stenocephala*) as well as resistance-associated SNPs. DESS is particularly valuable for remote sampling, where immediate refrigeration is not feasible and shipment on ice is prohibitive. Its compatibility with molecular diagnostics and ability to eliminate cold chain requirements make it an important tool for expanding parasitological research and surveillance in field settings.

## CRediT authorship contribution statement

**Yi-Jou Chen:** Investigation, Methodology, Validation, Data curation. **Vanessa Li:** Writing – review & editing, Investigation, Data curation. **Michelle Suwandy:** Investigation, Methodology, Data curation. **Ioana Bianca Mitrea:** Investigation, Methodology, Data curation. **Douglas Hayward:** Resources, Investigation, Data curation. **Susan Jaensch:** Resources, Investigation, Data curation. **Emily Kate Francis:** Writing – review & editing, Resources, Funding acquisition, Conceptualization. **Jan Šlapeta:** Writing – review & editing, Writing – original draft, Visualization, Validation, Supervision, Software, Resources, Project administration, Methodology, Investigation, Funding acquisition, Data curation, Conceptualization.

## Acknowledgements

Funding was provided by the Betty and Keith Cook Canine Research Bequest (Sydney School of Veterinary Science, The University of Sydney, Australia), and by the UEFISCDI - Executive Agency for Higher Education, Research, Development and Innovation Funding (Romania): PN-IV-P2-2.2-MC-2024 and 131 PED. This research was in part funded by the Sydney School of Veterinary Science Research & Enquiry Unit of Study 2025 fund. This work was completed in partial fulfillment for the requirements of the Doctor of Veterinary Medicine degree, The University of Sydney (VL).

## References

Abdullah, S., Stocker, T., Kang, H., Scott, I., Hayward, D., Jaensch, S., Ward, M.P., Jones, M.K., Kotze, A.C., Šlapeta, J., 2025. Widespread occurrence of benzimidazole resistance single nucleotide polymorphisms in the canine hookworm, *Ancylostoma caninum*, in Australia. Int. J. Parasitol. 55, 173–182. doi: 10.1016/j.ijpara.2024.12.001

Beknazarova, M., Millsteed, S., Robertson, G., Whiley, H., Ross, K., 2017. Validation of DESS as a DNA preservation method for the detection of *Strongyloides* spp. in canine feces. Int. J. Environ. Res. Public Health 14, 624. doi:

Bloomquist, R.F., Elston, D.M., 2024. What’s eating you? Hookworm and Cutaneous Larva Migrans. Cutis 114, E12–E15. doi: 10.12788/cutis.1136

Bolt, B.J., Rodgers, F.H., Shafie, M., Kersey, P.J., Berriman, M., Howe, K.L., 2018. Using WormBase ParaSite: An integrated platform for exploring helminth genomic data. Methods Mol. Biol. 1757, 471–491. doi: 10.1007/978-1-4939-7737-6_15

Burton, K.W., Hegarty, E., Couto, C.G., 2024. Retrospective analysis of canine fecal flotation and coproantigen immunoassay hookworm positive results in Greyhounds and other dog breeds. Vet. Parasitol. Reg. Stud. Reports 51, 101026. doi: 10.1016/j.vprsr.2024.101026

Callahan, B.J., McMurdie, P.J., Rosen, M.J., Han, A.W., Johnson, A.J., Holmes, S.P., 2016. DADA2: High-resolution sample inference from Illumina amplicon data. Nat. Methods 13, 581–583. doi: 10.1038/nmeth.3869

Dawson, M.N., Raskoff, K.A., Jacobs, D.K., 1998. Field preservation of marine invertebrate tissue for DNA analyses. Mol Mar Biol Biotechnol 7, 145–152. doi:

Evason, M.D., Weese, J.S., Polansky, B., Leutenegger, C.M., 2023. Emergence of canine hookworm treatment resistance: Novel detection of *Ancylostoma caninum* anthelmintic resistance markers by fecal PCR in 11 dogs from Canada. Am. J. Vet. Res. 84, 20230718. doi: 10.2460/ajvr.23.05.0116

Francis, E.K., Šlapeta, J., 2022. A new diagnostic approach to fast-track and increase the accessibility of gastrointestinal nematode identification from faeces: FECPAK(G2) egg nemabiome metabarcoding. Int J Parasitol 52, 331–342. doi: 10.1016/j.ijpara.2022.01.002

Gaither, M.R., Szabó, Z., Crepeau, M.W., Bird, C.E., Toonen, R.J., 2011. Preservation of corals in salt-saturated DMSO buffer is superior to ethanol for PCR experiments. Coral Reefs 30, 329–333. doi: 10.1007/s00338-010-0687-1

Gasser, R.B., Chilton, N.B., Hoste, H., Beveridge, I., 1993. Rapid sequencing of rDNA from single worms and eggs of parasitic helminths. Nucleic Acids Res. 21, 2525–2526. doi: 10.1093/nar/21.10.2525

Geary, T.G., Drake, J., Gilleard, J.S., Chelladurai, J.R.J.J., Jimenez Castro, P.D., Kaplan, R.M., Marsh, A.E., Reinemeyer, C.R., Verocai, G.G., 2025. Multiple anthelmintic drug resistance in the canine hookworm *Ancylostoma caninum*: AAVP position paper and research needs. Vet. Parasitol., 110536. doi: 10.1016/j.vetpar.2025.110536

Gonzálvez, M., Ruiz de Ybáñez, R., Rodríguez-Caro, R.C., Maíz-García, A., Gómez, L., Giménez, A., Graciá, E., 2022. Assessing DESS solution for the long-term preservation of nematodes from faecal samples. Res. Vet. Sci. 153, 45–48. doi: 10.1016/j.rvsc.2022.10.010

Harris, T.W., Arnaboldi, V., Cain, S., Chan, J., Chen, W.J., Cho, J., Davis, P., Gao, S., Grove, C.A., Kishore, R., Lee, R.Y.N., Muller, H.M., Nakamura, C., Nuin, P., Paulini, M., Raciti, D., Rodgers, F.H., Russell, M., Schindelman, G., Auken, K.V., Wang, Q., Williams, G., Wright, A.J., Yook, K., Howe, K.L., Schedl, T., Stein, L., Sternberg, P.W., 2020. WormBase: a modern Model Organism Information Resource. Nucleic Acids Res. 48, D762–D767. doi: 10.1093/nar/gkz920

Hawdon, J.M., Wise, K.A., 2021. *Ancylostoma caninum* and other canine hookworms, in: Strube, C., Mehlhorn, H. (Eds.), Dog Parasites Endangering Human Health. Springer International Publishing, Cham, pp. 147–193.

Howe, K.L., Bolt, B.J., Shafie, M., Kersey, P., Berriman, M., 2017. WormBase ParaSite - a comprehensive resource for helminth genomics. Mol Biochem Parasitol 215, 2–10. doi: 10.1016/j.molbiopara.2016.11.005

Jimenez Castro, P.D., Howell, S.B., Schaefer, J.J., Avramenko, R.W., Gilleard, J.S., Kaplan, R.M., 2019. Multiple drug resistance in the canine hookworm *Ancylostoma caninum*: an emerging threat? Parasit. Vectors 12, 576. doi: 10.1186/s13071-019-3828-6

Jimenez Castro, P.D., Willcox, J.L., Rochani, H., Richmond, H.L., Martinez, H.E., Lozoya, C.E., Savard, C., Leutenegger, C.M., 2025. Investigation of risk factors associated with *Ancylostoma* spp. infection and the benzimidazole F167Y resistance marker polymorphism in dogs from the United States. Int J Parasitol Drugs Drug Resist 27, 100584. doi: 10.1016/j.ijpddr.2025.100584

Kitchen, S., Ratnappan, R., Han, S., Leasure, C., Grill, E., Iqbal, Z., Granger, O., O’Halloran, D.M., Hawdon, J.M., 2019. Isolation and characterization of a naturally occurring multidrug-resistant strain of the canine hookworm, *Ancylostoma caninum*. Int. J. Parasitol. 49, 397–406. doi: 10.1016/j.ijpara.2018.12.004

Krauth, S.J., Coulibaly, J.T., Knopp, S., Traore, M., N’Goran, E.K., Utzinger, J., 2012. An in-depth analysis of a piece of shit: distribution of *Schistosoma mansoni* and hookworm eggs in human stool. PLoS Negl. Trop. Dis. 6, e1969. doi: 10.1371/journal.pntd.0001969

Leutenegger, C.M., Evason, M.D., Willcox, J.L., Rochani, H., Richmond, H.L., Meeks, C., Lozoya, C.E., Tereski, J., Loo, S., Mitchell, K., Andrews, J., Savard, C., 2024. Benzimidazole F167Y polymorphism in the canine hookworm, *Ancylostoma caninum*: Widespread geographic, seasonal, age, and breed distribution in United States and Canada dogs. Int. J. Parasitol. Drugs Drug Resist. 24, 100520. doi: 10.1016/j.ijpddr.2024.100520

Martin, M., 2011. Cutadapt removes adapter sequences from high-throughput sequencing reads. EMBnet.journal 17, 10–12. doi: 10.14806/ej.17.1.200

McKean, E.L., Grill, E., Choi, Y.J., Mitreva, M., O’Halloran, D.M., Hawdon, J.M., 2024. Altered larval activation response associated with multidrug resistance in the canine hookworm *Ancylostoma caninum*. Parasitology 151, 271–281. doi: 10.1017/S0031182023001385

Murali, A., Bhargava, A., Wright, E.S., 2018. IDTAXA: a novel approach for accurate taxonomic classification of microbiome sequences. Microbiome 6, 140. doi: 10.1186/s40168-018-0521-5

Oosting, T., Hilario, E., Wellenreuther, M., Ritchie, P.A., 2020. DNA degradation in fish: Practical solutions and guidelines to improve DNA preservation for genomic research. Ecol. Evol. 10, 8643–8651. doi: 10.1002/ece3.6558

R Core Team, 2022. R: A language and environment for statistical computing. R Foundation for Statistical Computing, Vienna, Austria. https://www.R-project.org/.

Rinaldi, L., Coles, G.C., Maurelli, M.P., Musella, V., Cringoli, G., 2011. Calibration and diagnostic accuracy of simple flotation, McMaster and FLOTAC for parasite egg counts in sheep. Vet Parasitol 177, 345–352. doi: 10.1016/j.vetpar.2010.12.010

Seutin, G., White, B.N., Boag, P.T., 1991. Preservation of avian blood and tissue samples for DNA analyses. Can. J. Zool. 69, 82–90. doi: 10.1139/z91-013

Sharpe, A., Barrios, S., Gayer, S., Allan-Perkins, E., Stein, D., Appiah-Madson, H.J., Falco, R., Distel, D.L., 2020. DESS deconstructed: Is EDTA solely responsible for protection of high molecular weight DNA in this common tissue preservative? PLoS One 15, e0237356. doi: 10.1371/journal.pone.0237356

Sonmez, H.I., Madak, E., Karaer, M.C., Sarimehmetoglu, H.O., 2025. Anthelmintic resistance in *Ancylostoma caninum*: A comprehensive review. Vet. Med. Sci. 11, e70434. doi: 10.1002/vms3.70434

Stocker, T., Scott, I., Šlapeta, J., 2023. Unambiguous identification of *Ancylostoma caninum* and *Uncinaria stenocephala* in Australian and New Zealand dogs from faecal samples. Aust. Vet. J. 101, 373–376. doi: 10.1111/avj.13272

Stocker, T., Šlapeta, J., 2025. Characterisation of β-tubulin isotypes in *Uncinaria stenocephala* and implications for benzimidazole resistance in hookworms. Vet. Parasitol. 339, 110569. doi: 10.1016/j.vetpar.2025.110569

Stocker, T., Ward, M.P., Šlapeta, J., 2024. Nationwide USA re-analysis of amplicon metabarcoding targeting β-tubulin isoform-1 reveals absence of benzimidazole resistant SNPs in *Ancylostoma braziliense*, *Ancylostoma tubaeforme* and *Uncinaria stenocephala*. Vet. Parasitol. 327, 110118. doi: 10.1016/j.vetpar.2024.110118

Venkatesan, A., Jimenez Castro, P.D., Morosetti, A., Horvath, H., Chen, R., Redman, E., Dunn, K., Collins, J.B., Fraser, J.S., Andersen, E.C., Kaplan, R.M., Gilleard, J.S., 2023. Molecular evidence of widespread benzimidazole drug resistance in *Ancylostoma caninum* from domestic dogs throughout the USA and discovery of a novel beta-tubulin benzimidazole resistance mutation. PLoS Pathog. 19, e1011146. doi: 10.1371/journal.ppat.1011146

Workentine, M.L., Chen, R., Zhu, S., Gavriliuc, S., Shaw, N., Rijke, J., Redman, E.M., Avramenko, R.W., Wit, J., Poissant, J., Gilleard, J.S., 2020. A database for ITS2 sequences from nematodes. BMC Genet. 21, 74. doi: 10.1186/s12863-020-00880-0

Yoder, M., De Ley, I.T., Wm King, I., Mundo-Ocampo, M., Mann, J., Blaxter, M., Poiras, L., De Ley, P., 2006. DESS: a versatile solution for preserving morphology and extractable DNA of nematodes. Nematology 8, 367–376. doi: 10.1163/156854106778493448

Zajac, A.M., Conboy, G.A., 2011. Veterinary Clinical Parasitology. Wiley.

